# An inexpensive setup for robust activity tracking in small animals: Portable Locomotion Activity Monitor (pLAM)

**DOI:** 10.1101/2021.08.21.457197

**Authors:** Yash Sondhi, Nicolas J. Jo, Britney Alpizar, Amanda Markee, Hailey E. Dansby, J. P. Currea, Samuel T. Fabian, Carlos Ruiz, Elina Barredo, Matthew Degennaro, Akito Y. Kawahara, Jamie C. Theobald

**Affiliations:** Department of Biology, Florida International University, Miami FL 33174 USA; Department of Psychology, Florida International University, Miami FL 33174 USA; McGuire Center for Lepidoptera and Biodiversity, Florida Museum of Natural History, University of Florida, Gainesville, FL 32611 USA; Imperial College, London, England

**Keywords:** activity detector, animal motion tracking, circadian rhythm, insect diel-activity, low-cost

## Abstract

1. Advances in computer vision and deep learning have automated animal behaviour studies that previously required tedious manual input. However, tracking activity of small and fast flying animals remains a hurdle, especially in a field setting with variable light conditions. Commercial locomotor activity monitors (LAMs) can be expensive, closed source, and generally limited to laboratory settings.
2. Here, we present a portable locomotion activity monitor (pLAM), a mobile activity detector to quantify small animal circadian activity. Our setup uses inexpensive components, is based on open-source motion tracking software, and is easy to assemble and use in the field. It runs off-grid, supports low-light tracking with infrared lights, and can implement arbitrary light cycle colours and brightnesses with programmable LEDs. We provide a user-friendly guide to assembling pLAM hardware and accessing its pre-configured software and guidelines for using it in other systems.
3. We benchmarked pLAM for insects under various lab and field conditions, then compared results to a commercial activity detector. They offer broadly similar activity measures, but our setup captures flight and bouts of motion that are often missed by beam-breaking activity detection.
4. pLAM will enable high-throughput quantification of small animal location and activity in a low-cost and accessible manner, crucial to studying behaviour that can help inform conservation and management decisions.

## Introduction

Improvements in computing speed, memory, and technology have helped automate animal behaviour studies that were once performed manually (M. W. Mathis & Mathis, 2020). For pose estimation, multi-animal recognition, and motion tracking, computers can sometimes exceed human accuracy (A. Mathis et al., 2018; Nath et al., 2019; Tadres & Louis, 2020). Unfortunately, many tools are inaccessible without sufficient computational power and programming proficiency (von Ziegler et al., 2021). Further, it remains challenging to track small or fast animals in the field, with variable light and weather. Although commercial solutions, like the Locomotor Activity Monitor (LAM, https://trikinetics.com/) and EthoVisionX (https://www.noldus.com/ethovision-xt), address some of these problems, they are often expensive and bulky (Table S1). Open-source solutions, when available, are specialized for model organisms, such as fruit flies or mice (Matikainen-Ankney et al., 2019)(Chiu et al., 2010), **(Matikainen-Ankney et al., 2019),** and generally limited to laboratory use. Field activity tracking, with camera traps and radio tags, are often only practical for large animals (Debata & Swain, 2018; Nunes-Silva et al., 2019; Pirie et al., 2016).

The lack of a portable and affordable tool has hindered behavioral data collection for smaller, non-laboratory animals, such as nocturnal arthropods, and only a handful of studies have attempted it (Table S2). They used either manual observation (Fullard & Napoleone, 2001), which is difficult to scale up, or non-portable setups (Edwards, 1960; J. L. Smith et al., 2016; P. H. Smith, 1983), which are difficult to replicate in the field. Light traps are easy to use and portable, but have inherent problems for inferring activity of phototactic animals (Lamarre et al., 2015; Lewis & Taylor, 1965). Because trap effectiveness decreases with distance, they bias towards animals present around a trap (Baker & Sadovy, 1978). Further, trap light can activate otherwise inactive animals, such as those waiting for dawn (Donahue, 1962), pers. obs. last author). Additionally, many animals are blinded by bright trap lights, and become inactive after they settle (Frank et al., 2006), making actual activity even more difficult to establish. Suction traps, that sample captured insects hourly, monitor activity without the problems associated with bright lights (Wright & Morton, 1995), but these are strongly affected by the spatial distribution of insect populations (Taylor & Carter, 1961), and require cumbersome manual sorting.

Many insects face growing extinction risks (Boyes et al., 2021; Janzen & Hallwachs, 2021; Wagner, 2020; Yang et al., 2021), yet with the exception of certain pest species (Lima et al., 2020), little attention has gone toward automated monitoring. Such systems are crucial to understanding and documenting baseline behaviors, especially as anthropogenic factors, such as light pollution, alter and stress the environment. Here, we address several limitations of previous methods by introducing the Portable Locomotion Activity Monitor (pLAM) to automate activity monitoring of small animals under arbitrary light conditions. We benchmark this equipment under different conditions, provide examples of field and laboratory use, and compare results with an existing commercial package. With low-cost, open documentation, and portability, anyone can monitor locomotion of multiple small animals simultaneously, making this a novel tool to reveal fine-scale, small animal activity patterns.

### Description and Implementation

The monitor logs activity events for an individual or group of animals in a small enclosure, detecting activity by comparing the difference in pixel values between successive images from a video stream. When pixel differences cross a predefined threshold, the program saves images or videos and logs the time and duration of the motion event. The pLAM can operate over a range of natural light conditions, or in the lab with programmable LEDs that simulate natural 24 light-dark cycles. It can run on power banks or battery packs in the field (Fig. 1,2, Table S3).

**Figure 1:**
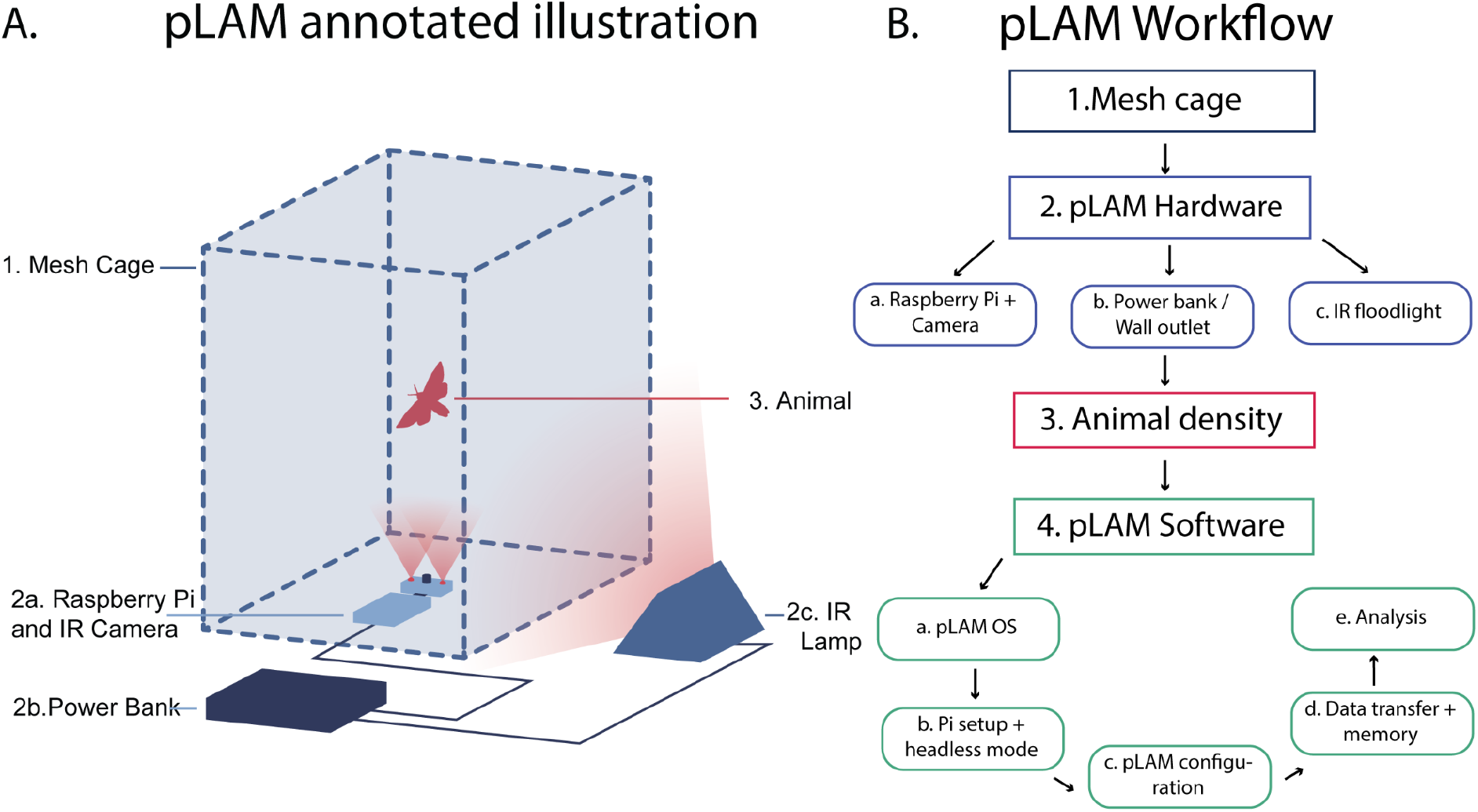
Annotated Illustration of pLAM A: 1. Mesh Cage, 2a. Raspberry pi and camera, 2b. Power bank, 2c IR floodlight 3. Small animal B: pLAM workflow (See Appendix 1-2 for more details)

**Figure 2:**
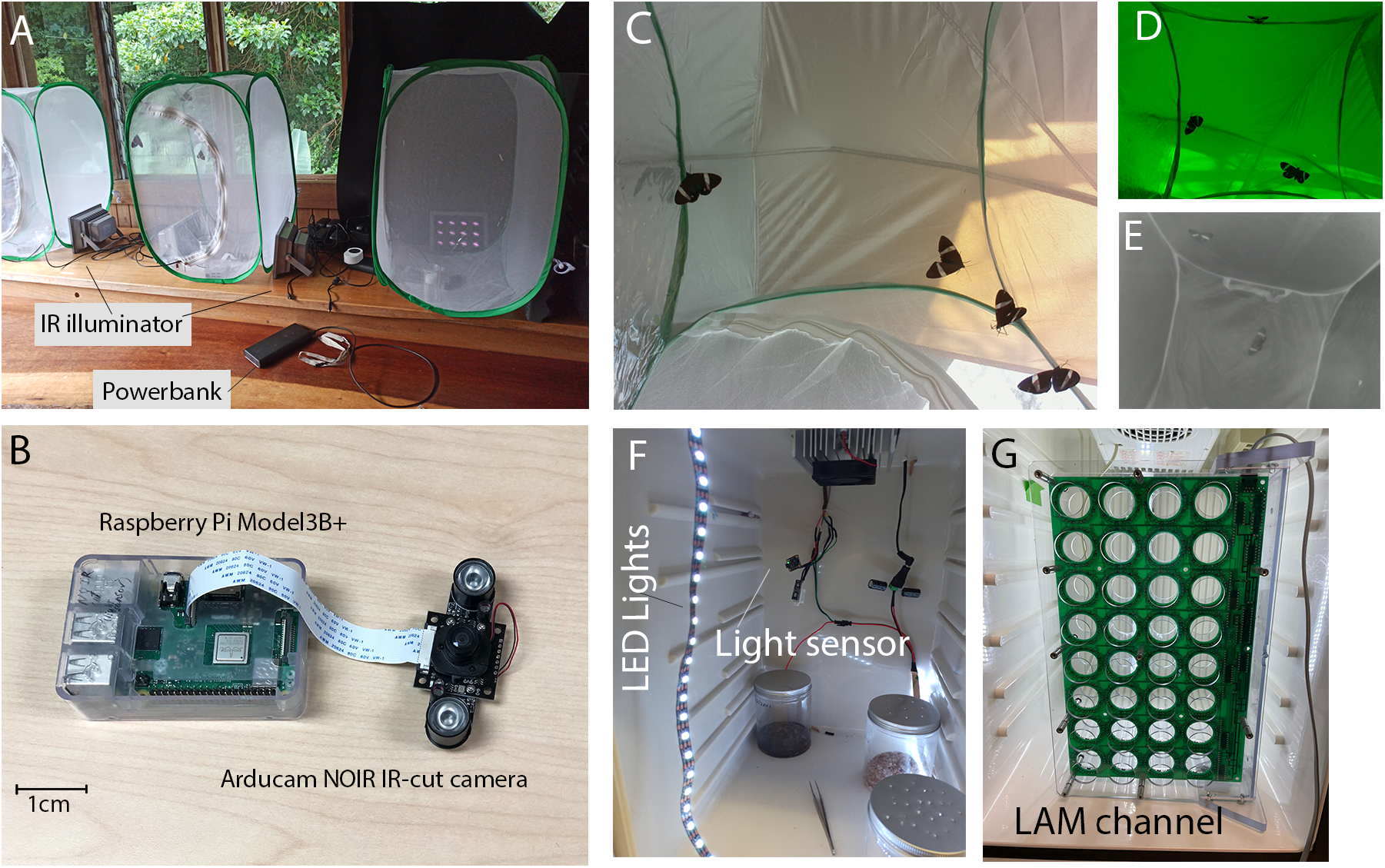
Different components of the activity chamber; A: Mesh tent containing raspberry pi and power bank. B: Raspberry pi and camera C-E: Image of multiple animals (moth:*Hypocrita excellens)* as seen by the pLAM under various light conditions C: Day view with visible light (IR-cut Filter) D: Day without IR cut filter, E: Night without IR filter F: Rearing Incubator with controllable lights and light sensor G: Commercial 32 channel LAM (TriKinetics) in modified incubator

pLAM consists of: (1) An infrared camera and visible lights, both controlled by a RaspberryPi single-board computer, and external infrared lighting for motion capture in the dark (Fig. 1,2, Table S3). (2) A command line interface for setting activity capture parameters. (3) Wrappers and pre-configured settings for running *motion* (https://github.com/Motion-Project/motion), an open-source library that detects motion between successive video frames. (4) A python-based processing pipeline to automate capture, logging, and analysis of activity with text files or images. (5) Scripts to control light settings in a light chamber of choice. We provide a step-by-step guide to building your own pLAM and using the associated software (Appendix 1). We also provide a detailed troubleshooting and optimisation guide to use the pLAM in different animals and conditions (Appendix 2). All code is available on github (www.github.com/yashsondhi/diel-light-pi) and the disk image of the pre-configured pLAM OS along with other instructables are hosted on OSF server (https://osf.io/8p5kw/).

## Methods

### Benchmarking

We benchmarked the pLAM with objects of different sizes and various test insects. We compared results to data from a commercial activity detector and tested our device with 15 insect species in the field.

a. **Object size and distance detection limit**: To test the detectable range of object size and distances, we suspended plastic beads in a chamber and moved them at random intervals with a fan. We then monitored diel-activity of three differently sized lab reared insects (see Supplementary Methods).
b. **Comparison with commercial activity detector LAM:** The LAM (Locomotor activity monitor) and DAM (Drosophila activity monitor) systems are commercially available activity monitors that are commonly used in the lab with small insects like fruit flies and mosquitoes. They are designed and distributed by Trikinetics (https://trikinetics.com/) and use infrared beams and detectors to monitor insect activity. Similar to a burglar detection system, an insect passing through a ring of infrared beams interrupts the signal to the detector, and the LAM device records this as a motion event (Fig. 2,<u>LAM25 Data Sheet</u>). Unlike the pLAM which monitors the collective activity of all individuals present in the field of view, LAMs monitor individual activity. They work well for small insects in the lab, but are expensive, with setup costs between 1000-4000 USD, and cannot house larger animals or function in natural settings. We generated comparative reference diel activity data with the LAMs for the species used in the pLAM trials (Fig. 3, Supplementary Methods).
c. **Field tests:** To benchmark the pLAM in the field, we conducted trials in Monteverde, Costa Rica. We set up 6 pLAMs outdoors at the edge of a cloud forest reserve. We collected different insect species from light traps, then monitored activity with the pLAMs. We used 4-10 individuals of each species for up to 48 hours under natural illumination and environmental conditions (Fig. S4, Supplementary Methods).

**Figure 3:**
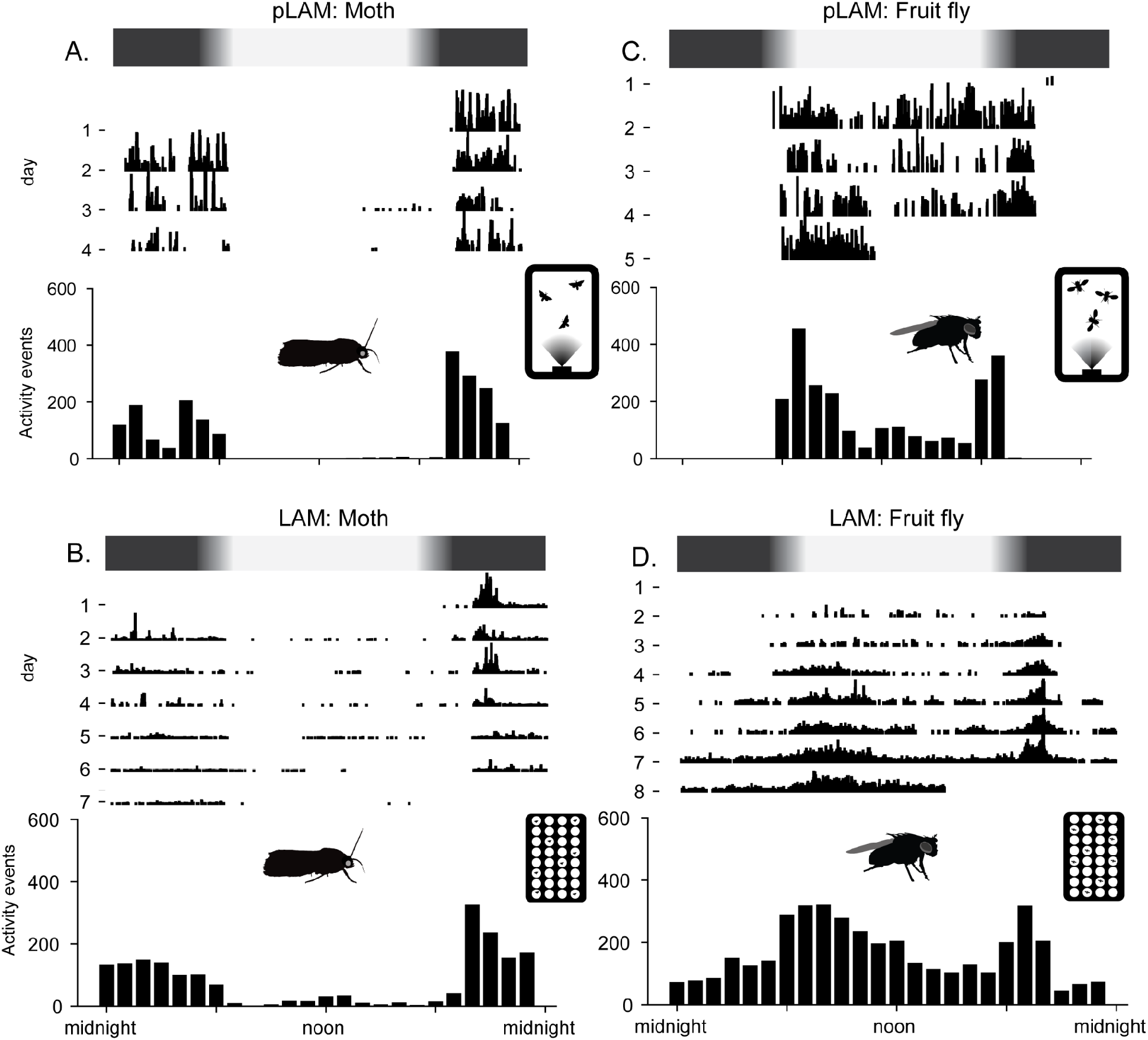
Comparison of pLAM data with LAM data for two different sized animals. A: pLAM data for Wax moth (*Galleria mellonela*, body size ^∼^10 mm); B: LAM data for *G*. *mellonela* C-D: C: pLAM data for Fruit fly (*Drosophila melanogaster*, body size ^∼^3 mm); D: LAM data for Fruit fly (*D*.*melanogaster*). Black and white bar at the top of each graph represents the light cycle.

### Data analysis

**pLAM**: Our software logged motion events in a text file and saved corresponding images of frames with maximal pixel differences. For bead benchmark trials, we examined motion event frequency as density functions. For insect activity trials, we measured the scaled pixel difference (normalised to the maximum pixel difference detected) of each image of every motion event, across all the days of the trial (Fig. 3A,C), and summary data, representing hourly frequency counts of motion events (Fig. 3A,C). Field-trials (Fig. S2) generated noisier data, and we therefore only included trials where activity patterns were consistent across both days without manual filtering of background motion. **LAM**: LAMs count beam breaking events in each channel per minute. We combined channel data and compared scaled motion event counts (Fig. 3 B,D). **Light**: We measured illuminance (lux) over the trial period (Fig. S3), also represented as a light bar above the plots.

## Results

### Object size and distance detection limit

To test the range of pLAM object detection, we suspended beads of different sizes at selected distances from the camera, and generated erratic motion with wind from a fan. Flat lines would represent no motion, but the monitor detected events for all conditions (Fig. S1). Large beads generated longer motion bouts, and small beads generated high frequency, short duration events. The smallest detectable beads were 4mm at over 60 cm away. We then proceeded to test various insects (wax moths ∼10mm, mosquitos ∼3mm and fruit flies ∼2mm) with an artificial, graded light-dark cycle in lab conditions. The device worked for all three, monitoring 4-5 days of activity (Fig. 3 A,C, Fig. S2).

### Comparison with commercial activity detector LAM

To test the accuracy of the pLAM activity data, we compared the same three species in standard commercial LAM activity detectors. Wax moths showed nocturnal activity with a peak in the first few hours of the night, and an overall pattern consistent across the LAM and the pLAM (Fig. 3 A,B) setups. Fruit flies were mostly diurnal and displayed peaks of activity at dawn and dusk. The pLAMs recorded mosquito activity exclusively during the day, peaking near dusk, but the LAM showed a symmetrical activity at dawn and dusk. (Fig. S5). The pLAMs recorded less baseline fruit fly activity than the LAMs (Fig. 3 C,D, Fig. S5). Images revealed pLAMs exclusively counted bouts of flight, rather than walking, which could be changed by altering the camera angle.

### Field tests with multiple species

To ensure devices were robust for field-use, we tested them at Estación Biológica Monteverde (EBM) in Costa Rica (see Supplementary Methods, Fig S4). We monitored 15 species (Fig. S4. see Table S4) in natural light and weather conditions, demonstrating pLAMs functioned well in the field, but are prone to signal noise, and require software and hardware modifications not necessary in the lab (Appendix 1). We found large variations in diel activity of species collected at light screens, contradicting suggestions that nocturnal collection indicates natural nocturnal activity.

## Discussion

The portable locomotion activity Monitor (pLAM) can track collective and individual small animal activity using frame-based difference methods for detecting motion under various light conditions (Fig. 1,2). We have evaluated its performance under lab and field conditions, and provide a detailed guide to hardware and software, and offer logistical advice for using it outdoors. (Appendix 1) We discuss our rationale for choosing pLAM components, compare it with other options, and discuss its limitations (Table S4). We detail the issues we had with lab and field trials and provide some general recommendations for using the pLAM in both conditions (Appendix 2).

### Rig design

We tested several camera and light combinations (Appendix 1-2) but eventually used (1) the raspberry NOIR cameras modified by Arducam with IR cut filters, for their wide field of view and function under both bright and dim, infrared lit, scenes. (2) IR LED illuminators, which are inexpensive and sufficiently light for moderately sized cages. (3) Raspberry Pi 3’s. Newer models increase cost but add little to data collection performance (although they do offer faster transfer speeds and USB3 ports). All components together typically cost $130-$180 USD (Table S3), roughly 10-fold cheaper than commercial options.

### Comparison of pLAM with other methods and commercial tools

Aside from their price, several commercial tools offer capabilities comparable to pLAM (Table S1). Options vary in interface, coding environment, and methodology but they come into two main categories useful for research. First, commercial software and hardware marketed towards tracking animal activity using either IR beams or video and can be used for circadian rhythm monitoring. These are often expensive and can require a subscription. Second, open-source programs which, although free, rarely work in field conditions without modifications to the code and equipment and can rely on contrasting backgrounds or ample lighting. The pLAM offers an open-source solution that functions in the field with inexpensive hardware. It does have limitations, the false positives are high if the camera detects background motion from trees, wind or humans, and it may fail to capture the entire field of view.

### Guidelines for recording diel-activity with pLAM

pLAM works consistently in the lab with controllable lighting. Isolating the setup in temperature-controlled incubators is ideal, but at minimum, shield the pLAM from external light and motion. The Pi supports ambient light and temperature with extra sensors, but commercial wireless loggers are more convenient. Troubleshooting the pLAM and lights is easier without live animals, although further optimisation with test animals is usually required to getting robust data. For mosquitos, flies and moths, we found when keeping humidity levels high and providing a food source, we got data for 5-7 days post eclosion. With small cages, 20-25 flies or mosquitoes were needed to get an accurate representation of the activity, but with medium cages for the moths, 4-8 individuals were enough. It is possible to record even a single individual, but since the animal occasionally leaves the field of view, the activity data could be incomplete. We limited the insect motion to smaller chambers entirely within the camera’s field of view, but this reduces the animal’s tendency to fly. We noticed for flies and mosquitos, the pLAM was more likely to pick up flying behaviour than walking behaviour, but this was not an issue for the larger moths. While setting up pLAM experiments in different systems, we recommend altering animal density, cage size, using higher resolution cameras, lowering the motion threshold and using stronger IR illumination to get more robust data.

Conducting pLAM trials in the field is slightly more challenging. Access to electrical mains, internet and shelter from wind and rain and light pollution are ideal, but often hard to obtain. Portable routers, UPSes and lightweight tents offer relatively inexpensive solutions, however even they fail to withstand prolonged harsh weather. Severe weather causes power fluctuations, random background motion of the forest and water leakage on the electrical equipment and should be avoided. We had two setups get blown away during the field trials and eventually used a large indoor library with windows to provide access to light while shielding the setups from wind.

### Variation in diel-activity of insects in the field

Although this study was meant to be a test of pLAM in the field, we were surprised at the diel-activity patterns we found across various species. Despite using only species that were attracted to light traps, which have often been used as an indicator of being nocturnal (Akite et al., 2015), we found tremendous variation in their diel activity. Some were completely nocturnal, but some were active throughout the day and night and others showed peaks of activity at dawn and dusk. This data suggests that a much more detailed and systematic survey of diel-activity is required across small animals especially nocturnal species.

In conclusion, we plan to optimize the device, add more features, and improve the software. We learnt from our failures and plan modifications that will improve the robustness of the setup in field-conditions including implementing machine learning filters post-data collection. We hope that this tool and the accompanying guide to building and using your own pLAM will help promote field-based studies of diel-activity periods and eventually lead to the creation of a large database of diel-activity periods across animal taxa.

## Acknowledgements

We thank Chandra Earl, Emily Ellis, and Scott Cinel for testing and prototyping pLAM. Caroline Storer, Nick Homzick, David Plotkin, Lilian Hendricks and Sarah Steele Cabrera provided valuable discussion and feedback. Michael Ramone helped with mosquito care and experiments. The Estación Biológia de Monteverde, Costa Rica provided laboratory space to allow the field component of research to be conducted.

## Funding

Financial support was provided by NSF IOS 1920895 to AYK, and NSF BCS-1525371 to JPC and NSF IOS-1750833 to JCT, YS received support from the Presidential Fellowship, a Tropical Conservation Grant and National Geographic Explorer Grant and a Lewis Clark Exploration Grant from the American Philosophical Society.

## Author Contributions

YS: Conceptualisation, writing,
NJJ: Data collection, analysis, writing
BA: Data collection, writing
AM: Data collection, Rig testing
STF: Writing+illustrations
CR: Rig design, writing
EB: Data Collection and writing
MD: Conceptualisation, funding.
JPC: Rig design, Data Collection, Data Curation, Data analysis
CT: Rig design
*AYK: Conceptualisation, supervision, writing, funding
*JCT: Conceptualisation, supervision, writing, funding, data analyses

## Supplementary Information

*Appendix 1: Step by step instructable*

https://www.dropbox.com/s/wxldf7h1h2wvnn6/Instructable%20for%20activity%20chamber_2021_07_01_shared.docx?dl=0

*Appendix 2: Guide to troubleshooting and setting up the various components:*

https://www.dropbox.com/s/b0zvndsggrrne3p/Troubleshooting%20and%20optimisation%20Guide.docx?dl=0

*Supplemental methods*

https://www.dropbox.com/s/n92kl9iumn39jev/Supplemental%20Methods.docx?dl=0

## Supplementary Figures

**Figure S1:**
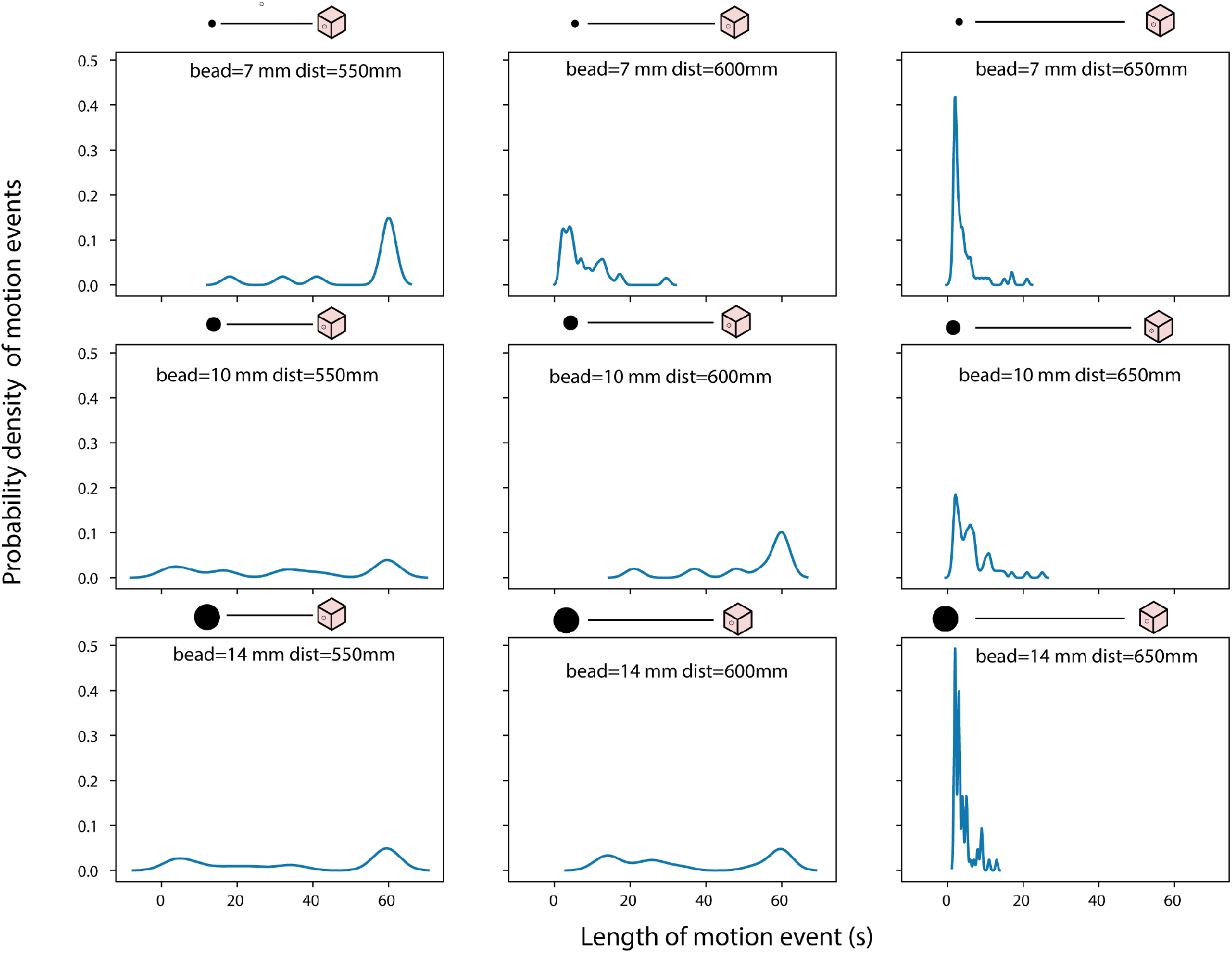
Benchmarking with beads as proxies for insects using different bead sizes and lengths from the detector. The graphs depict the probability density function of the length of different motion events.

**Figure S2:**
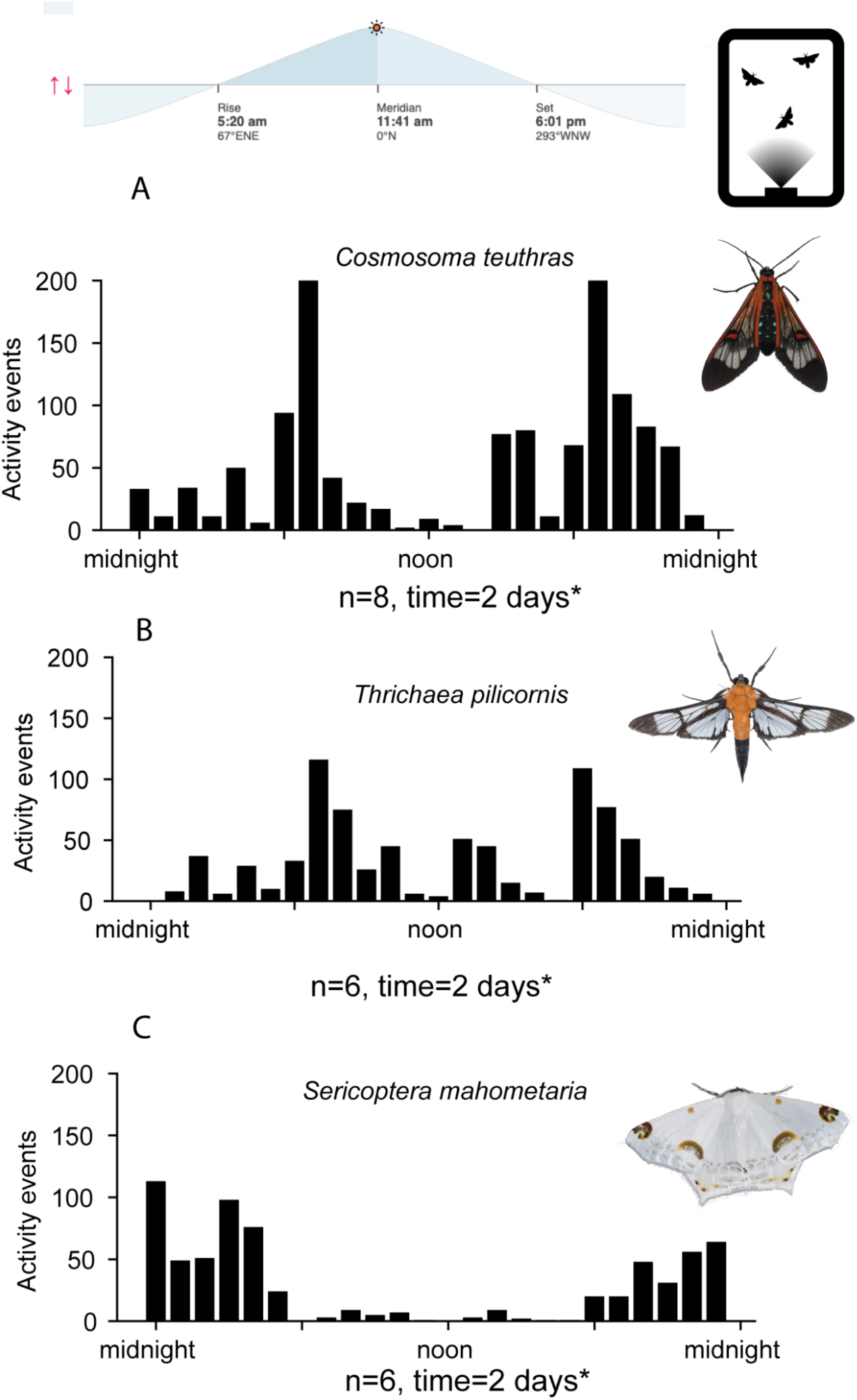
Data from field trials A-C, A: Erebidae: Arctiinae, *Cosmosoma teuthras*, n=8, t=2 days. B: Crambidae: *Trichaea pilicornis*, n =6, t = 2 days. C: Geometridae: *Sericoptera mahometaria*, n = 6, t= 2 days. * no data for 4 hours during the afternoon on day 2 of trial due to a power failure. n = number of individuals tested in one trial, time= duration of trial.

**Figure S3:**
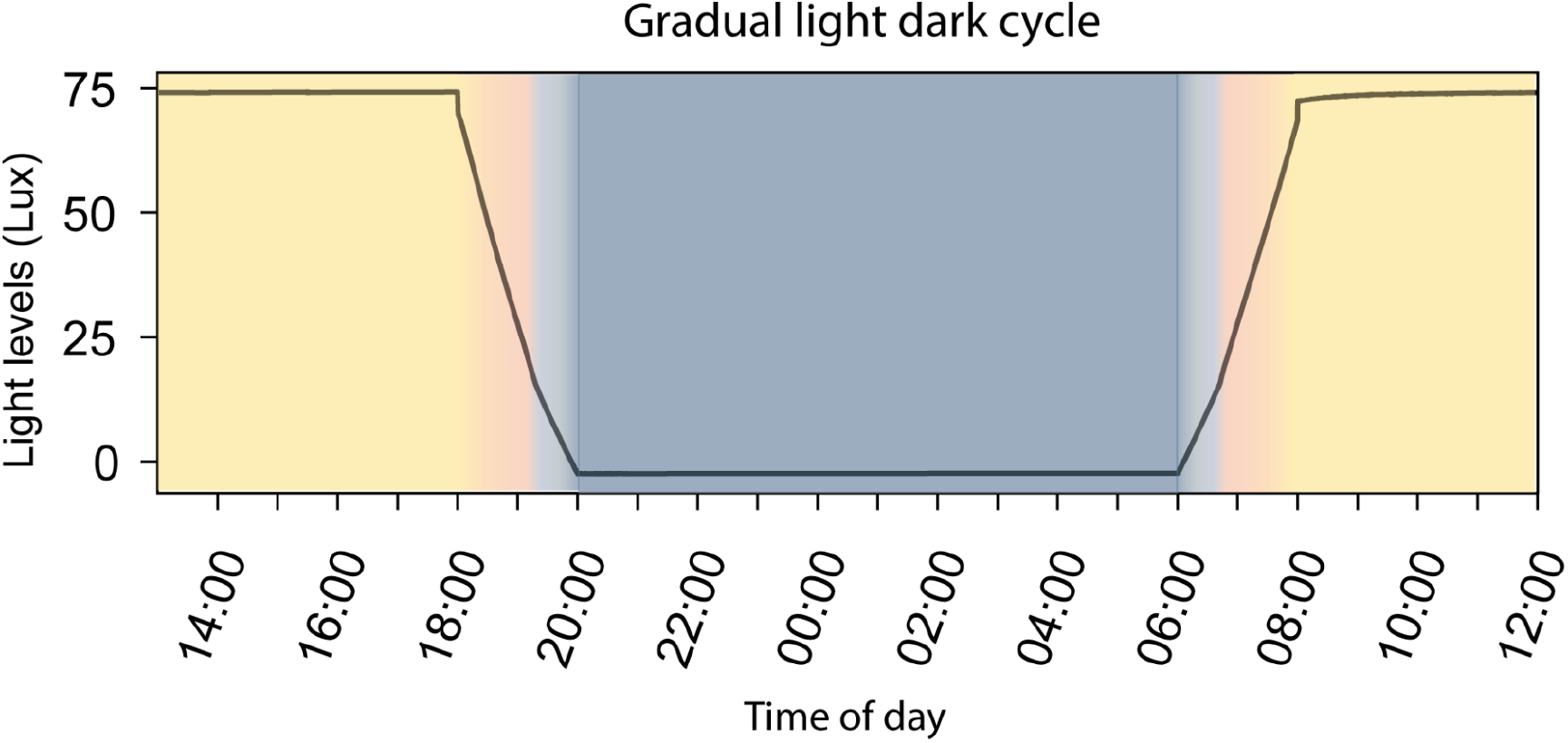
Gradual light dark cycle used in the lab for the pLAM and the LAMs. The absolute intensity differed for different chambers, but the relative intensity was the same. The measurements were obtained using a Adafruit TSL5291 sensor.

**Figure S4:**
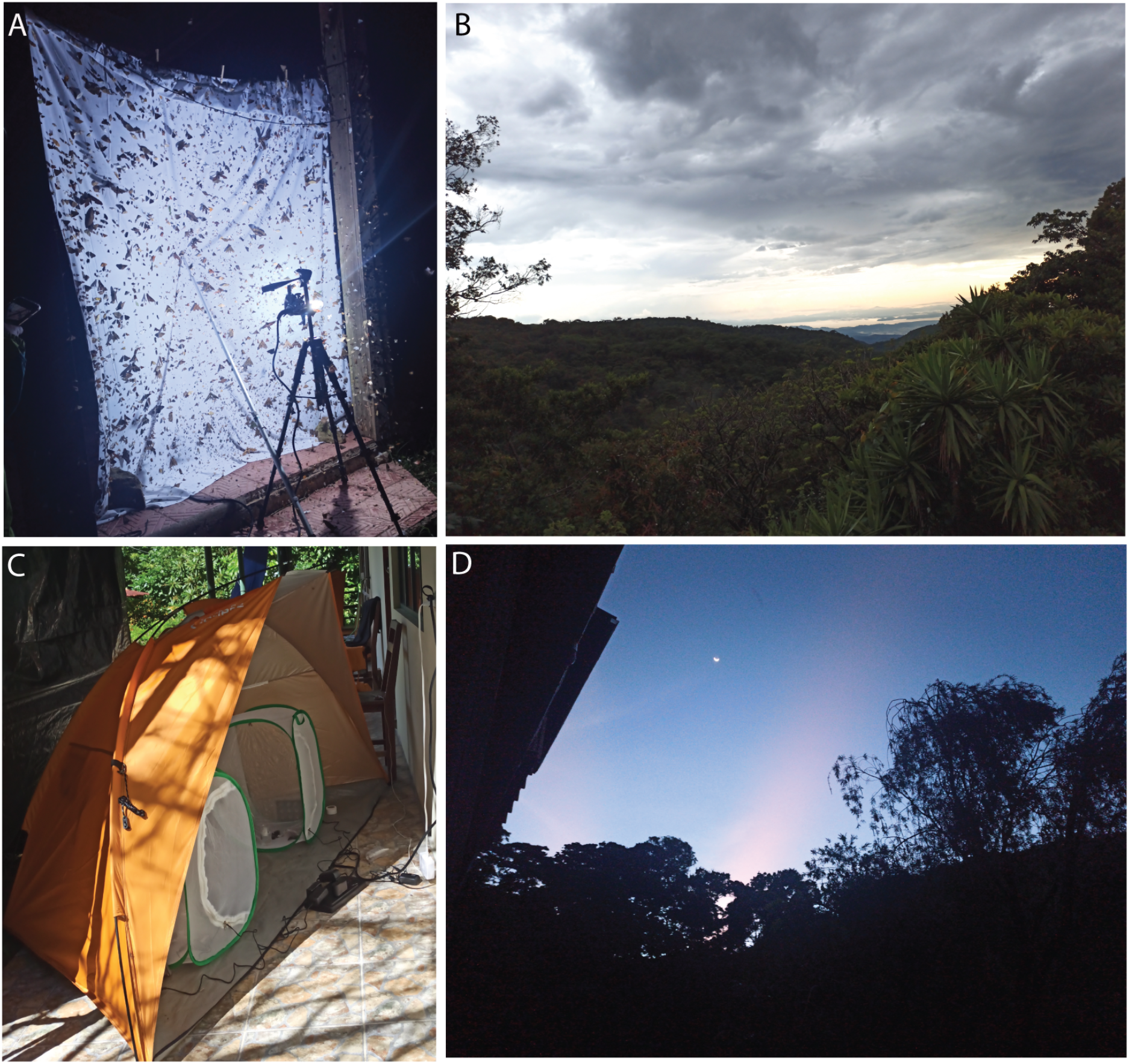
Field-based images of a light sheet, habitat and pLAM setup. A: Metal halide light in front of at white sheet at Estación Biológica Monteverde (EBM), Costa Rica, B: Habitat around EBM C: pLAM in tents at the field-station. D: View of the sky at dawn from EBM

**Figure S5:**
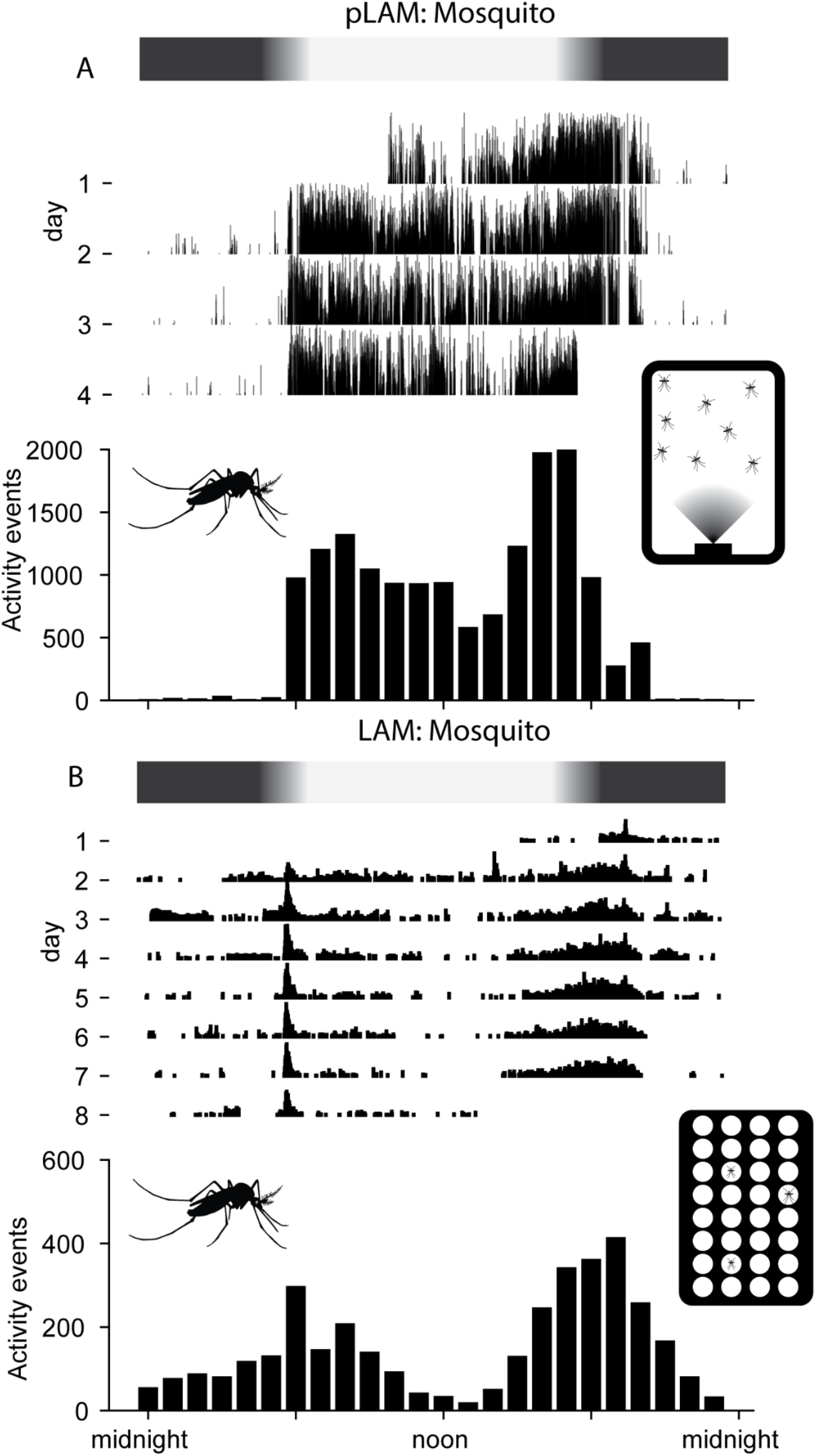
Comparison of pLAM and LAM data for male mosquitoes (*Aedes aegypti*). A: pLAMs with 20-25 male *A. aegypti.* B: LAMs with 32 channels with A. *aegypti*, data shown is only from males (n=16) over time= 8 days.

**Supplementary Table S1:** Other available software that monitors circadian rhythm with information about their cost, interface, user environment, tracking method and comments about their utility.

| Software Name | Open Source | Interface | Env. | Method | Comments | Link |
| --- | --- | --- | --- | --- | --- | --- |
| TriKinetics | No<br>Hardware can be \$400 to upwards of \$3000 | GUI | App | IR beam arrays | Commonly used for recording insect locomotion in lab-reared insects. Works with LAM and DAM hardware | https://trikinetics.com/ |
| Big Brother | No<br>\$40 a channel and paid software | GUI | App | Video camera recording movement | Uses time-based distance travelled and can be used to track circadian rhythm, not as useful for flying organisms | https://actimetrics.com/products/big-brother/ |
| "MouseActivity" | Yes | Command-based | MATLAB | Video camera | Uses a tablet to record video. Similar to pLAM, but is more stringent and seems to require better contrast with the background as it converts inverted grayscale image to binary and uses that to analyze after video is recorded. Optimised for walking behaviour with larger organisms and not applicable for outdoor field trials. | https://bmcresnotes.biomedcentral.com/articles/10.1186/s13104-020-4916-6 |
| "Tracker" | Yes; Not on github but on Brandeis website for free | Command-based | Java | Video camera mixed with IR beams | Similar methodology to pLAM by using text-based coordinates and images at 1 second intervals to record movement without recording a video. However, it uses a reference image to compare and requires a white background to contrast the dark flies and so it cannot be applied to the outdoor field trials. Useful for recording long trials with low volume of data. | https://journals.plos.org/plosone/article?id=10.1371/journal.pone.0037250 |
| ZooTracer | Yes | Command-based | OpenCV | Analysis pre-recorded video | Not directly used for circadian rhythm but it uses a pre-recorded video to analyze movement of unmarked individuals. Useful for the field but it can't detect what kind of animal is interacting and isn't useful for limiting storage usage | https://www.microsoft.com/en-us/research/project/zootracer/?from=http%3A%2F%2Fresearch.microsoft.com%2Fen-us%2Fprojects%2Fzootracer%2F |
| EthoVision XT | No. \$2000/year or \$6000 for a one-time license | GUI | N/A | Either pre-recorded video or live feed | Really expensive but has nice features including calibration features, setting field in view, and extras for specific species. Def not worth price tag | https://www.noldus.com/ethovision-xt |
| Behavior Cloud | No. \$990/year | App iOS/mobile or mac/PC | N/A | Either pre-recorded video or live feed | Analyzes data for you including movement tracking, coding complex behaviors, and even heart rate/EKG | https://www.behaviorcloud.com/ |
| Track 3D Insects | Price available on request (estimated cost 10-20k) | App | N/A | Records video to analyze | Tracks movement over time but it does track coordinates in 3D by using 2 separate cameras and even uses IR for night experiments. Custom solutions are designed for each lab. | https://www.noldus.com/track3d/insects |
| Graphite | Yes | GUI | App | Video recordings of human readable tags | Not directly used for diel-activity monitoring, but could be adapted for diel-activity monitoring if individual identity is | https://besjournals.onlinelibrary.wiley.com/doi/10.1111/2041- |
|  |  |  |  |  | required | 210X.13200 |

**Supplementary Table S2:**
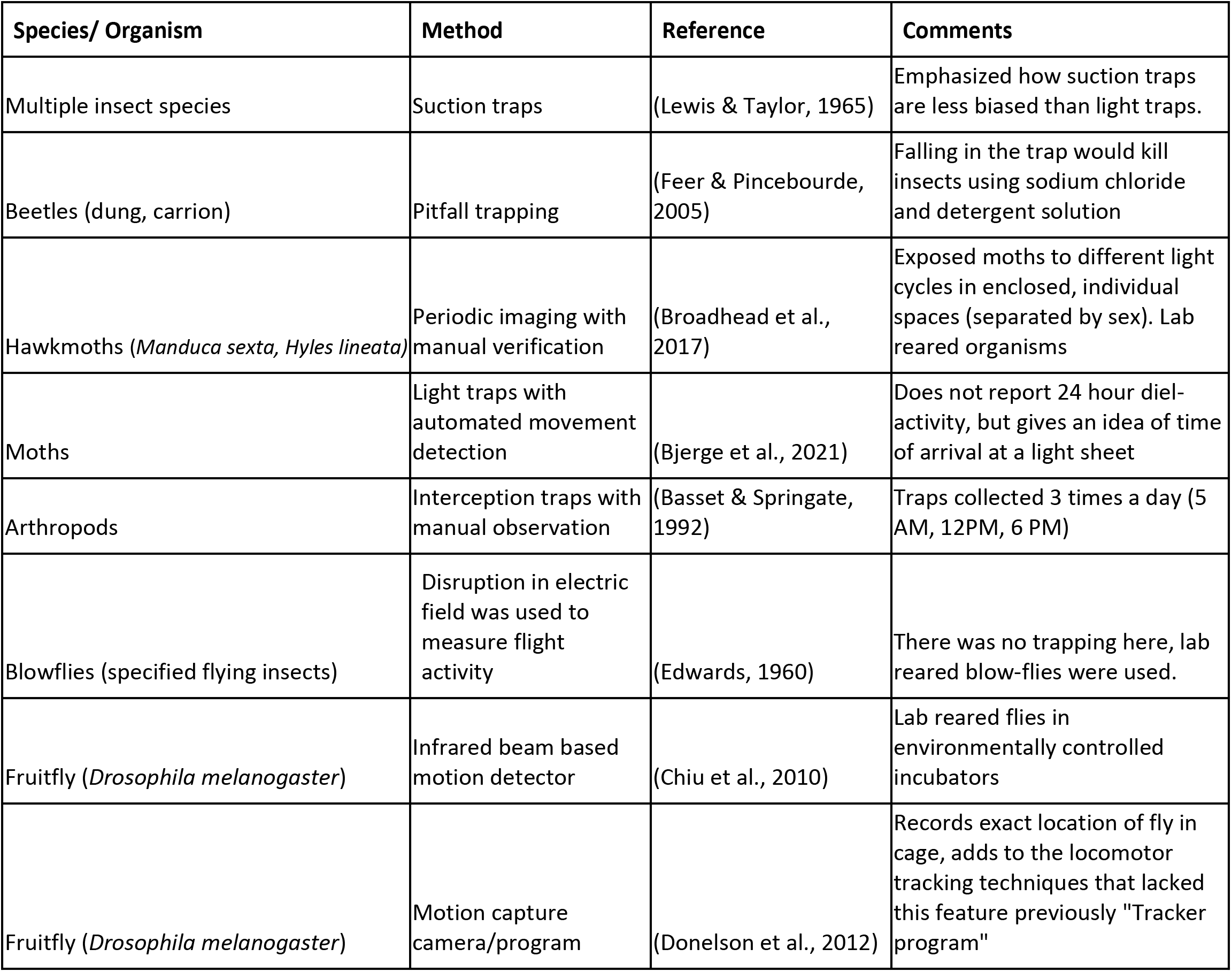
Studies that monitored small animal (arthropod) diel-activity, the species and methods they used and comments about the study.

**Supplementary Table S3:** Components, potential suppliers, and approximate cost in US dollars to build a single pLAM.

| Sr. | Part name | Cost (USD) | Supplier | Comments |
| --- | --- | --- | --- | --- |
| no. |  |  |  |  |
| 1 | Raspberry pi model 3+ kit | \$55 | CanaKit via amazon | Can buy the Pi separately for (\$35) and buy power supply and case separately |
| 2 | Adafruit NOIR camera V1 (5 MP) | \$30 | ArduCam via amazon | Can buy the 8 MP NOIR camera, but without the IR-cut feature images in the day are strangely coloured |
| 3 | Power bank/ Portable UPS | \$30-45\$ | Inui via amazon/Cyber power via amazon | A 20000 mAh power bank lasts approximately 12 hours. For longer use utilise wattage power banks/ portable UPS (Cyber power 750 VA). Alternatively connect to mains via low wattage UPS PiHat, CyberPower 325VA |
| 4 | Memory card | \$12-\$24 | Sandisk via amazon | 64GB or 128 GB are recommended depending on length of recordings required |
| 5 | Extra IR lights (optional) | \$30 | JC illuminator via amazon | Different brand of IR illuminators can work, use ones that turn-on automatically in low light |
| | | | | Total=\$127-\$184 |

**Supplementary Table S4:**
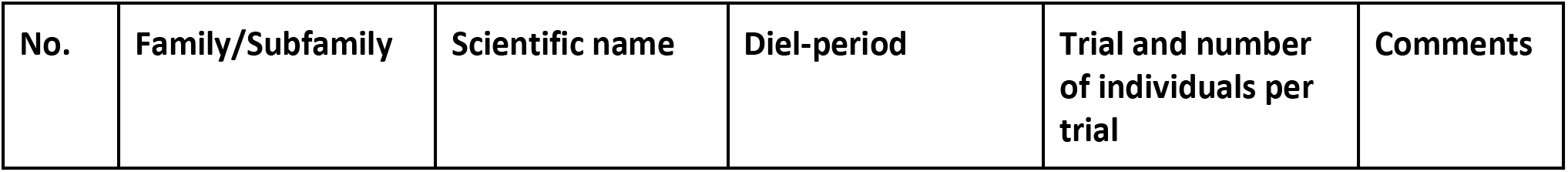
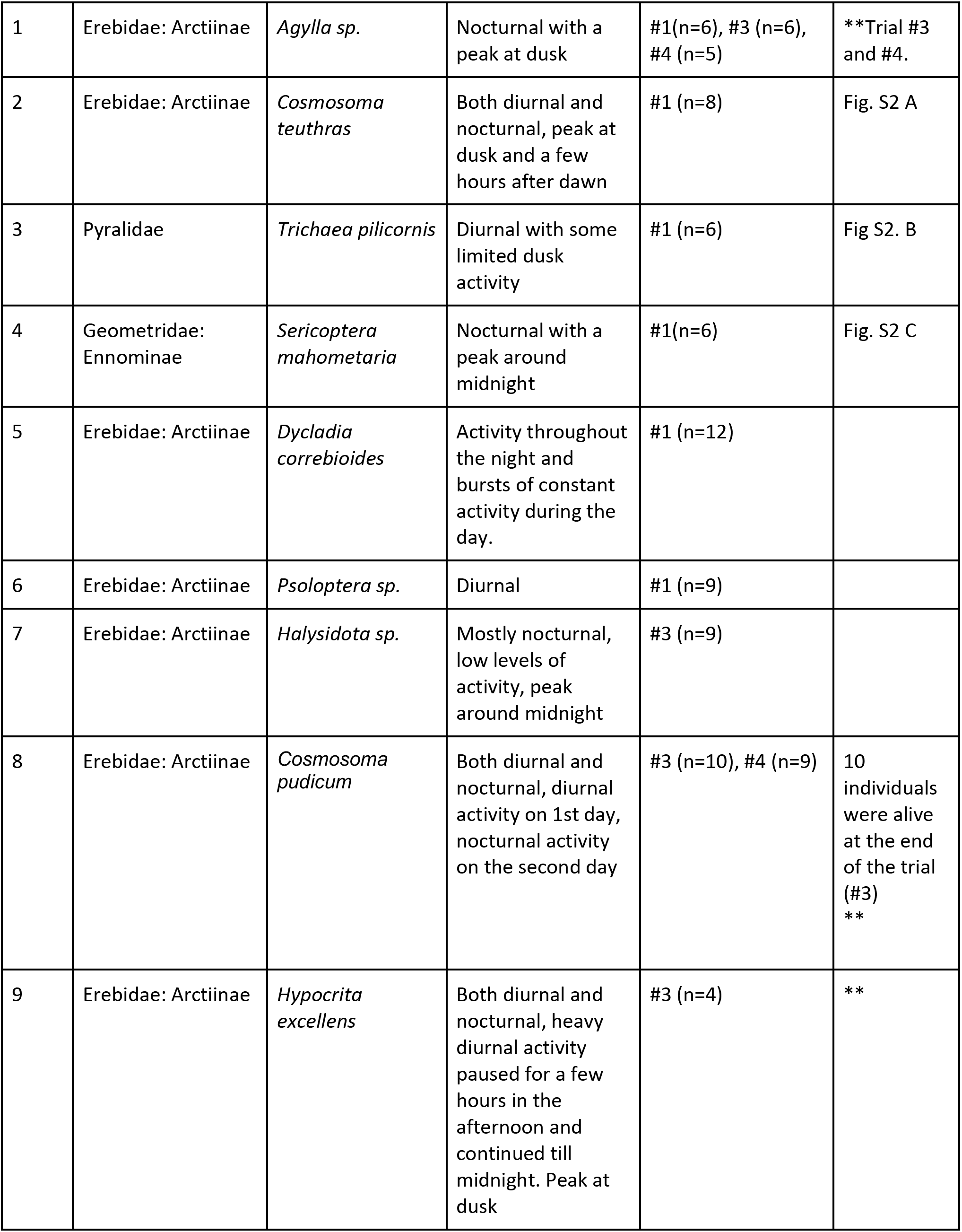
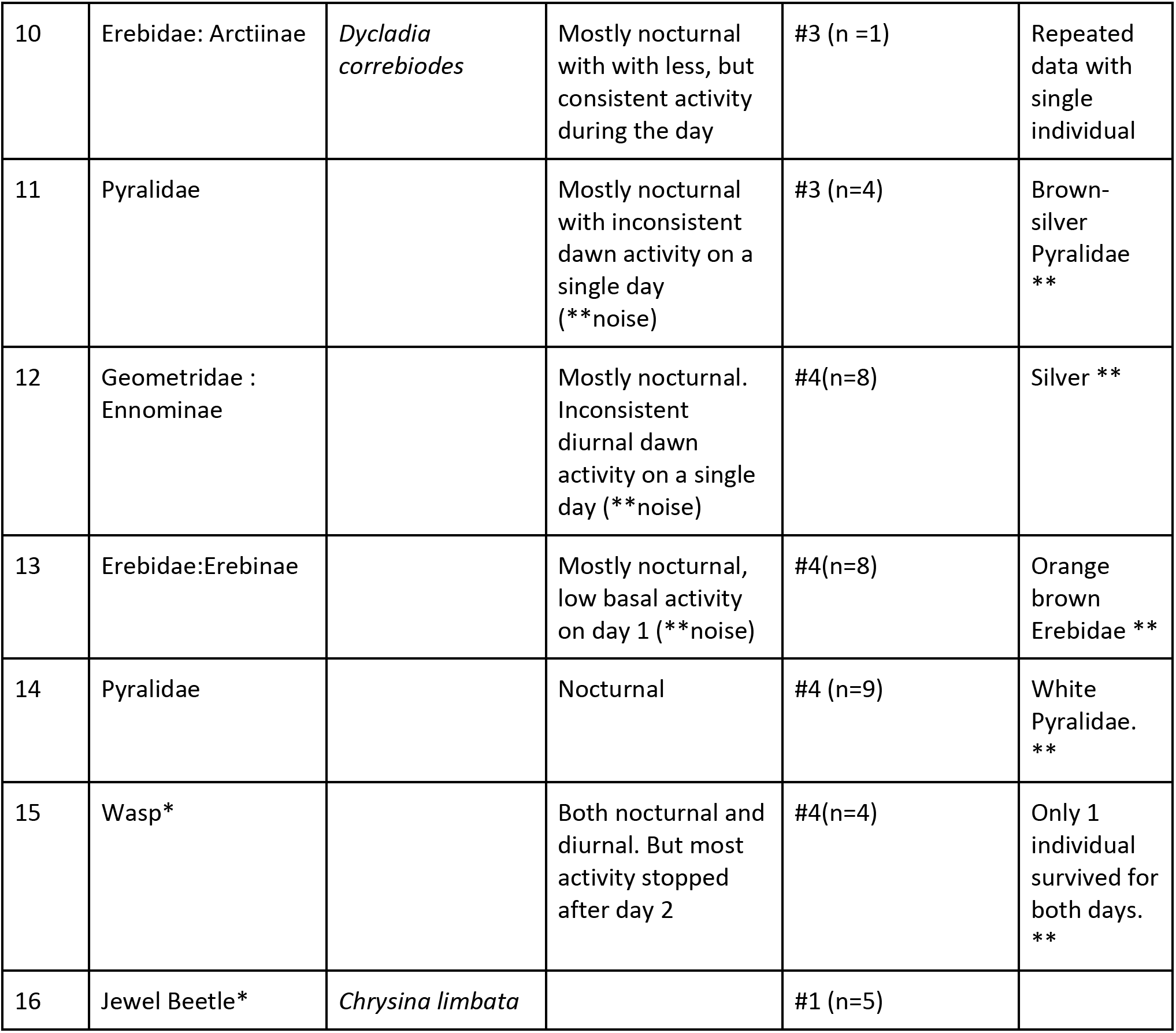
Species tested in the field pLAM trials in Monteverde Costa Rica. * Unless mentioned otherwise, all species tested were moths. ** Had motion activity from the trees and requires additional filtering.

## Notes

https://github.com/yashsondhi/diel-light-pi

https://osf.io/8p5kw/

### Competing Interest Statement

The authors have declared no competing interest.

