## Appendix 1 for "An inexpensive setup for robust activity tracking in small animals: Portable Locomotion Activity Monitor (pLAM)"

Tutorial for setting up an insect 24 hour activity detector

Version 4: Updated 20 August 2021

Code is available at: <https://github.com/yashsondhi/diel-light-pi/>

For more detailed tutorials and guides refer to [https://osf.io/8p5kw/]

**Overview**

This tutorial has instructions to build an activity detector that allows monitoring insect activity over a 24 hour window. The activity detector can track when an insect in a small cage is moving. It uses a Raspberry Pi (single board computer) and open source motion capture tools (motion) to monitor and detect insect motion in a small chamber. By adding infrared lights and using a small infrared-camera it is possible to track insects throughout the night. The same camera switches to the non-infrared mode during the day to ensure monitoring in all light conditions. In addition the raspberry pi can control visible lights using controllable LEDs

**Introduction:**

Part 1: Setup the Raspberry Pi and motion

Part 2: Assemble and attach the NOIR camera and IR lighting

Part 3: Run motion using custom scripts and test motion-detection

Part 4: Test your device with an insect

Part 5: Analyse results and tweak parameters

Part 6: Optional : Make it portable and field-ready

Part 7: Optional: Setting up dynamically controllable lights

**Part 1: Setup of Raspberry Pi**

What do I need?

- Raspberry Pi 3 B+ (B Plus)
- Power supply 2.5A Micro USB Power Supply
- 64/ 128 GB micro-sd cards.
- Memory card reader/ micro-sd-card to SD card adapter
- HDMI cable + Monitor4
- USB Keyboard*
- USB mouse*
- Disk image file with custom OS
- Software to write disk image to SD-card (E.g. for mac users AppleBaker v2)

Optional extras

- A Raspberry Pi-Case
- Heat sinks

*Wi-fi and Bluetooth peripherals work too, but they need to be compatible with Linux

Use the instructions below to set-up a custom SD Card with the software

A working version of the activity detector software and operating system is available as a downloadable file. This file is a custom *image file* and needs to be written (copied) to an SD card using a *disk writing program*.

Available Raspberry Pi 3: Jesse: (https://osf.io/8p5kw/)

**V1.2 : Data collection and analysis software**

Mac users

- If you are using a Mac, install the Apple Pi baker V2 software to clone the SD card from the disk image (Link supplied).
- <https://www.tweaking4all.com/software/macosx-software/applepi-baker-v2/>
- Download the *image file* to your computer. You will get a zip file.
- Insert the memory card into the adapter/SD-card reader and connect it to your computer.
- Choose the 64 gb/ 128 gb memory card after left clicking select disk.
- Click Restore – And select the downloaded file

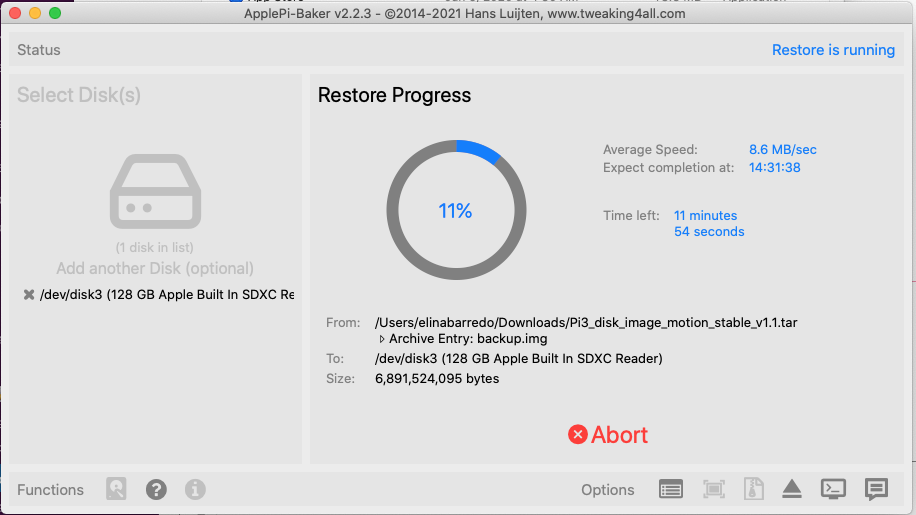

- Wait till process finishes (*it may take several minutes or even hours*). If it works you will get a boot drive show up on your desktop

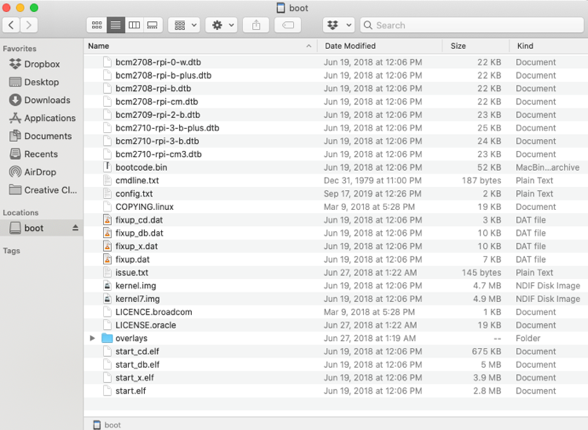

- Eject the boot drive on your computer and disconnect the SD card
- Insert the SD-card into the Raspberry Pi and power it on

**Windows/ Linux**

Use balena Etcher for Windows or Linux (<https://www.balena.io/etcher/>) or Raspberry Pi Imager. You may have to unzip the files.

After the file has downloaded, proceed to set up the raspberry pi using the tutorial below.

(<https://projects.raspberrypi.org/en/projects/raspberry-pi-setting-up/1>)

Using and setting-up your Raspberry Pi

- Connect your Pi
- Continue with connecting the Pi to a keyboard, mouse and monitor. See <https://projects.raspberrypi.org/en/projects/raspberry-pi-setting-up/3>

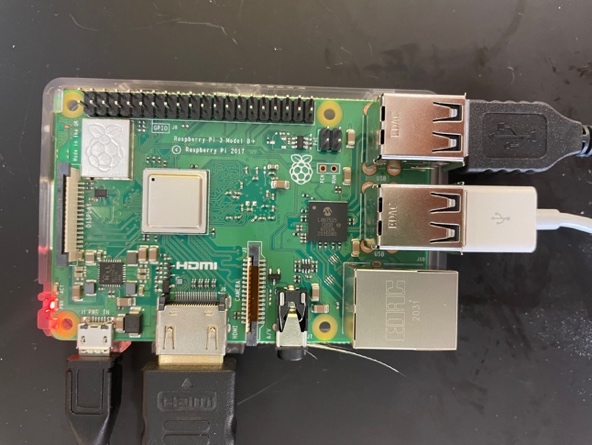

- Start your Pi (<https://projects.raspberrypi.org/en/projects/raspberry-pi-setting-up/4>)
- Finish Set up (<https://projects.raspberrypi.org/en/projects/raspberry-pi-setting-up/5>)
- Expand the file system. Expand the file system using (<https://geek-university.com/raspberry-pi/expand-raspbian-filesystem/>)
- Note: Type “sudo raspi-config” on your terminal and run to get configuration mode. Choose “Advanced options” in order to “Expand the file system”.

**Part 2: Assemble and attach the NOIR camera and IR lighting**

What do I need

- Day-Night Vision for Raspberry Pi Camera**^1^**, Arducam All-Day Image All-Model Support, IR LED for Low Light and Night Vision, M12 Lens Interchangeable, IR Filter Switch Programmable, OV5647 5MP 1080P

Optional extras

- Brighter IR Light LED Panel
- Default Brightness : Two small IR sensors attached to the Arducam NOIR camera
- Moderate Brightness : 20-30 USD LED IR panel, several are available on amazon. Can be found using the key words: Wide Angle IR Illuminator for Night
- High Brightness : Larsen electronics : IR floodlight LED IR Illuminator Light - 20 LED - 60 Watts - 9-42V - 750/850/940nm - 750'L X 110'W(-Flood-750nm-Black)
- Pi-Cam extension cable
- Follow the instructions and the tutorial to setup the camera and test it
- Attach heat sinks to the back of the two IR lights of the NOIR Arducam.
- Connect the NOIR camera^1^ to the raspberry pi after removing all the peripherals as shown below.
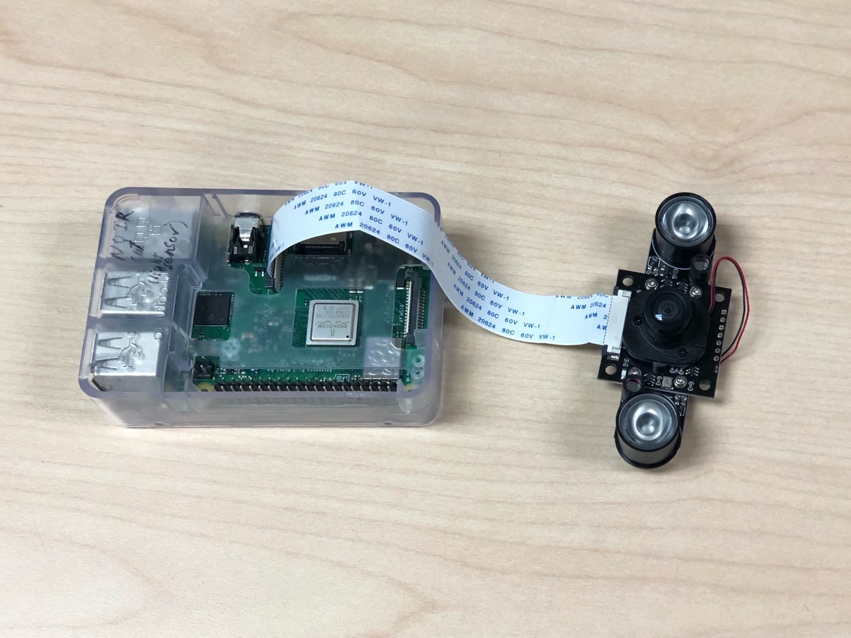

(<https://maker.pro/raspberry-pi/tutorial/how-to-interface-pi-noir-v2-camera-with-raspberry-pi>)

- Remove the lens cap
- Run **“raspistill -o test.jpg”**
- Set up IR lighting if required for recording night activity. The default Arducam in the materials has two IR LED’s that work well for troubleshooting and testing, but for brighter lighting, use IR LED panels. Refer to IR lighting at the end.

**^1^** You can modify the set-up to use different cameras but then you will have to reconfigure motion to work, also if you are not using the custom rom, refer to the motion documentation on installing the dependencies for pi-cams / webcams for your version Raspberry Pi OS.

**Part 3: Run motion using custom scripts**

What do I need?

- Custom image file installed on the raspberry pi (previously done in **Part 1**)
- Basic knowledge of Linux command line including changing files and folders using cd and running pre-written python scripts (<https://www.raspberrypi.org/documentation/linux/usage/commands.md>)

Follow the instructions below and test if motion works.

- Open the raspberry Pi
- Open a terminal (the black and blue box on the top left)
- Ensure you are connected to the internet
- If a *diel-light-pi* folder exists delete it and install the updated version from GitHub <https://github.com/yashsondhi/diel-light-pi>
- Install the code using the commands git clone <https://github.com/yashsondhi/diel-light-pi.git>
- Before running a motion trial, modify the file *project.conf* in the *configs* folder. Modify the output file name, organism, project name and user and for best results use motion_best.conf or motion_best_small.conf.
- Run “**python3 run_diel-light.py --setup**” and select *Yes* to continue
- Run “**python3 run_diel-light.py --run”** and select *Yes* to continue
- Wave your hand in front of the camera or smile.
- Press Ctrl+C (this stops the run) and look in the folder and the logs file to see if any motion events were detected (photos will pop-up)
- Test the hardware as well if all else fails
- To stop running use Ctrl+C *

**Part 4: Test an insect**

What do I need?

- Large Monarch Butterfly Habitat, Giant Collapsible Insect Mesh Cage Terrarium Pop-up 24 x 24 x 36 inches or any transparent cage of choice
- Raspberry pi camera with the IR cam and extra IR lights if required
- Any insect

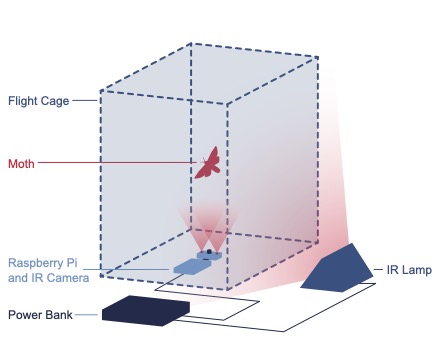

Test motion sensor on insect

- Place the IR camera and raspberry Pi in the bottom of the cage as shown in the figure above.
- Connect the power, monitor and peripherals
- Place the insect in the cage and close
- Turn on IR light if required for monitoring in the dark
- Confirm the parameters in the config file. Run “**python3 run_diel-light.py --run”**
- Wait……
- If you know the animal has moved stop (Ctrl+C) and look for the output folder
- Alternatively disturb the insect and run “**python3 run_diel-light.py --run”** again
- Wait…. Repeat process to see if the device works

**Part 5:** **Analyse Results and tweak parameters**

Check the output using analysis scripts and personalize motion configuration parameters
What do I need?

- A text editor and quick read through the motion configuration file and the analysis configuration file
- Patience
- pLAM OS v1.2 (to plot data on the pi)

The script generates log files for each trial, which is saved as a *text file* in each folder, this analysis script uses these log files and plots the data

**Analyse output**

- Open the raspberry Pi
- Open and modify the file “*configs/analysis.conf*” using a text editor.
  Change the following:
- **INPUT_PATH***: Path to the working directory which contains the log files of the data to be analysed
- **TREATMENT***: the word or pattern present in the log file name of the experiments you want to analyse. Data is extracted from all the log files with treatment in the filename.
- Save a copy of the *analysis.conf* file in the folder where you want to analyse data and note the file path
- For example *pi/home/diel-light-pi/2021_07_06-pi04/analysis.conf*

**Analyse data**

- Open the terminal
- Navigate to *diel-light-pi/scripts*
- Make sure that *analysis_pipeline.py* is present
- Run the script **“python3 analysis_pipeline.py --conf** [replace_with_the full_path_to_analysis.conf]” –extract (e.g. “**python3 analysis_pipeline.py –conf pi/home/diel-light-pi/2021_07_06-pi04/analysis.conf –extract”**)
- Look for a *csv* file in the output folder and check to see if the motion events were saved
- If it looks fine run the script “**python3 analysis_pipeline.py** –**conf** [replace_with_your_path_to_analysis.conf] **--plot_test**”
- This plots raw pixel difference values across all days, a histogram of the motion events and the average or median value of pixel difference for each hour. It is quick to plot and a good check of the data.
- For more complete publishable plots that take longer to plot, use command “**python3 analysis_pipeline.py –conf** [replace_with_your_path_to_analysis.conf] --**plot_all**”
- This plots scaled pixel-difference and summary plots of hourly counts of motion events.

*You must change these values to the folder with the data for the script to work, other values are recommended but not required since the script will run with the incorrect names.

Parameters that can be modified optionally:

- **OUTFILENAME**: text string, Descriptive name used for output files and plot headers
- **CSVFILE**: File name of combined csv
- **USEFILTER**: True or False, if pixel difference values have very high points, set filter = True
- **FILTER_MAX**: Integer value, maximum value of pixel difference value to use. Example, 10000 (do not use commas)
- **FILTER_MIN**: Integer value, minimum value of pixel difference value to use, For example, 100
- **YLIM**: Integer value, Y limit of summary count graph, set to upper bound of values present (usually between 200-600)

Repeat (**Step 4)** and tweak different parameters till you are happy with results

- Try a larger insect if you get motion to work with your hand, but not the insect. You may also increase the number of insects tested.
- If you get images but not all the time, tweak the sensitivity of the device by changing the motion configuration file.(<https://motion-project.github.io/motion_config.html>)
- Play around with the [threshold] number of pixels that change to qualify as motion, it can be anywhere between 100-200 to the maximum number of pixels in the image size you specify.
- Change the image size : Smaller images need fewer pixels to detect motion and can have faster frame-rates, but are bad at detecting small objects.
- Add more light
- You can refer to the user forum of motion as well as the internet to see what else you can modify. Motion has many configuration options including masking certain regions, taking videos instead of images and noise control, video generation, and using multiple cameras as well as triggering a set of scripts on detecting motion. It can be extended in numerous ways. Scripts to convert time images to videos are also available upon request.
- There are some sample configuration files available in motion like *motion_snapshot.conf* (time-lapse), *motion_best.conf* (best image per motion event, i.e. image with the most motion) and *motion_best_small.conf* (same as *motion_best* but with a lower threshold).

**Part 6: Optional: Make it portable and field-ready**

What do I need

- Portable Power bank recommended >10,000 mAH
- Portable Home UPS/ Portable generator for powering IR lighting
- Transparent Tent/ Waterproof tarpaulin
- Access to internet to ensure system time is updates (optional)
- Access to a screen+ keyboard and mouse to troubleshoot the pi **OR**
- Wi-Fi hotspot + VNC viewer to access the raspberry pi headless.

I recommend using a portable wi-fi hotspot and setting up VNC viewer that allows you to use a laptop to control your raspberry pi when it is running headless.
You can also use a LAN cable and try connecting to it using to SSH. ssh pi@raspberrypi.local after SSH is enabled (though you may need a screen and monitor to enable SSH once) (<https://www.dexterindustries.com/howto/connecting-raspberry-pi-without-monitor-beginners/>). Use the command **hostname -I** to get the ethernet IP address which can be used to connect to the GUI via VNC.

**Set-up the raspberry Pi and detector to be portable**

- This can be achieved in many ways. I recommend buying a portable screen and keyboard with a trackpad and a long HDMI cable. This works fairly well in the field and requires less technical troubleshooting.
- Access to an internet connection is recommended as this ensures the raspberry pi can update its time, but it is not required, and the alternate is to manually force the time to change at the beginning of the experiment
- Connect the Pi to the power bank and add the Bluetooth keyboard and the monitor with the long HDMI
- Alternatively get the Pi to run headless and start connected to a fixed Wi-Fi on boot^1^, by changing the priority of the Wi-Fi connection you want it to connect to.
  (<https://www.raspberrypi.org/documentation/configuration/wireless/headless.md>). You can also install VNC viewer and connect to the PI using a phone or laptop^2^. (<https://magpi.raspberrypi.org/articles/vnc-raspberry-pi>) or (<https://www.raspberrypi.org/documentation/remote-access/vnc/>)
- Add the IR lights and heat sinks if required to the PI
- Run your setup and get data
- Ensure the setup is housed in some sort of waterproof tent or casing if rain is a concern
- Running a pi headless with VNC viewer and a static IP address(<https://www.circuitbasics.com/how-to-set-up-a-static-ip-on-the-raspberry-pi/>) makes it easier to run in a cage and only requires the PI to be run of a power bank. The lights do not have to be in the cage and can be run of large portable power sources or mains

^1^This method is recommended only for advanced users, it is convenient but it does require more patience, troubleshooting and technical skill to get running.

^2^This method does not work well if you are on an encrypted Wi-Fi or are in a place where you cannot ensure your pi connects to a known network.

**Part 7: Optional: Setting up dynamically controllable lights**

**Overview:** Use single addressable lights to change the brightness of each LED. The raspberry pi can program each light individually and set the desired brightness.

What do I need?

- NeoPixel Adafruit lights (<https://learn.adafruit.com/adafruit-neopixel-uberguide/basic-connections>).
- If you have issues wiring it to the pi read more here (<https://learn.adafruit.com/neopixels-on-raspberry-pi/overview>) docs:
- And check this for setting up the software <https://circuitpython.readthedocs.io/projects/neopixel/en/latest/>
- 5V 2A power supply
- 1000 uf capacitor
- 470 micro ohm resistor
- Use the following tutorials to setup the lights (<https://magpi.raspberrypi.org/articles/neopixels-python>)
- Once the lights are functioning run the “**sudo python3.8 diel-light-pi/scripts/smooth_light_control.py --test** ”.
- To see the parameters of the light cycle use “**sudo python3.8 diel-light-pi/scripts/smooth_light_control.py --setup**”
- If you want to modify the parameters, edit the value in the script using a text editor.
- On rebooting the script should run by default and the lights should turn on and cycle if they are connected
- To verify the light readings, you can optionally use a TS2591 light sensor (Raspberry Pi light sensor TSL2591/ TSL2561 (<https://www.adafruit.com/product/1980> ) to check the readings using python save_light_data.py

**Troubleshooting**
Check the **“/etc/rc.local**” file and check that the path to the script is correct

| **Problem** | **Troubleshooting** |
| --- | --- |
| Failure to save photos or “camera not found” error | Check the camera connection. The ArduCam may be lose and not being read by the raspberry pi. |
| Timezone (clock) is wrong | Check the internet connection. Try your cell phone hotspot if Wi-fi is unreliable to synchronise the raspberry-pi  OR  Change time manually if internet is not available |
| The Pi and other gadgets shut off unexpectedly or seem to overheat | Check the voltage capacity of your power supply and any adaptors you are using. This may be more common if LEDs are also connected to the same multi outlet adaptor |
| The Pi runs out of space prematurely | You may have failed to “Expand the file system”  Go to **sudo raspi-config** |
| The SD card crashes completely | You can reuse the same SD card by going through the “Restore” steps from **Part 1**. Make sure to “Expand the file system” once you insert the card back into the pi. |

- !Advanced users only (ignore if you downloaded the disk image with the software)
- Alternatively, if you are already an advanced user, you can install the operating system yourself, or use an existing raspberry pi and install the motion capture tools For more instructions on installing noobs from scratch and setting up motion:
- Setting up Noobs (<https://www.raspberrypi.org/documentation/installation/installing-images/README.md>)
- Setting up Motion (<https://motion-project.github.io/motion_config.html>)
- Guide on how to set up a web-cam server (<https://www.instructables.com/id/How-to-Make-Raspberry-Pi-Webcam-Server-and-Stream-/>)
- For headless start up follow these guides: <https://howchoo.com/pi/how-to-set-up-raspberry-pi-without-keyboard-monitor-mouse>
- For Installing motion on Buster and later: <https://www.nickjvturner.com/blog/2019/10/05/setup-motion-from-scratch>
- Python versions <3.8 may not be compatible with new light sensor and humidity sensor software so please update to use them. https://installvirtual.com/how-to-install-python-3-8-on-raspberry-pi-raspbian/

**Amazon links**

Raspberry Pi 3 B+

<https://www.amazon.com/CanaKit-Raspberry-Power-Supply-Listed/dp/B07BC6WH7V/ref=sr_1_5?dchild=1&keywords=raspberry+pi+3b+plus+kit&qid=1627404464&sr=8-5>

Arducam NOIR Camera

<https://www.amazon.com/Arducam-Day-Night-Raspberry-Automatic-Switching/dp/B07X1VGQSL>

micro SD-Card

<https://www.amazon.com/Silicon-Power-Superior-MicroSD-Adapter/dp/B07S5QW5V3/?tag=androidcentralb-20&ascsubtag=UUacUdUnU69352YYwYg>

<https://www.amazon.com/SanDisk-microSD-Memory-Adapter-MICROSD-ADAPTER/dp/B0047WZOOO>

IR floodlight

<https://www.amazon.com/Infrared-Illuminator-High-Power-Analogue-Surveillance/dp/B01MYNZ7TY/ref=asc_df_B01MYNZ7TY/?tag=hyprod-20&linkCode=df0&hvadid=216554715043&hvpos=1o1&hvnetw=g&hvrand=17842431196575373743&hvpone=&hvptwo=&hvqmt=&hvdev=c&hvdvcmdl=&hvlocint=&hvlocphy=1015033&hvtargid=aud-801738734305:pla-352964030839&psc=1>

IR LED

<https://www.amazon.com/gp/product/B07DWQZLJF/ref=ppx_yo_dt_b_search_asin_title?ie=UTF8&psc=1>

Power Bank

<https://www.amazon.com/s?k=iniu+power+bank+20000mah&gclid=Cj0KCQjw3f6HBhDHARIsAD_i3D8KFWMvqwjEoUBLSoKIHiiWCUJuUnWjmFJRZBQ3m8pH4gwUOwAvd5MaAqHcEALw_wcB&hvadid=511486610291&hvdev=c&hvlocphy=9053030&hvnetw=g&hvqmt=e&hvrand=17036255343428179221&hvtargid=kwd-881805372682&hydadcr=5024_10680615&tag=googhydr-20&ref=pd_sl_4iw0yt548_e>

UPS Power Strip 350VA

<https://www.amazon.com/CyberPower-CP350SLG-Standby-Outlets-Compact/dp/B004OR0V2C/ref=asc_df_B004OR0V2C/?tag=hyprod-20&linkCode=df0&hvadid=167126093426&hvpos=&hvnetw=g&hvrand=6674934528120388502&hvpone=&hvptwo=&hvqmt=&hvdev=c&hvdvcmdl=&hvlocint=&hvlocphy=9053030&hvtargid=pla-312494979732&psc=1>

Controllable lights

<https://www.amazon.com/ALITOVE-Individual-Addressable-Programmable-Non-Waterproof/dp/B01MG49QKD?pd_rd_w=L7KNu&pf_rd_p=ed2d9dd9-5e5e-4b68-81ad-abff01eb6452&pf_rd_r=EGSZDHE76HHNRSJR53YW&pd_rd_r=6e9bb88d-410f-4f60-87a0-5ac1c6e16426&pd_rd_wg=2sHp5&pd_rd_i=B01MG49QKD&ref_=pd_bap_d_rp_1_i&qty=3&th=1&psc=1&selectObb=new>

5V Power supply for LEDs

<https://www.amazon.com/SHNITPWR-Converter-Transformer-5-5x2-5mm-Controller/dp/B07Q26YG61/ref=sr_1_28?dchild=1&keywords=5v+power+supply&qid=1614710818&s=electronics&sr=1-28>

1000 uF capacitors

<https://www.amazon.com/5pcs-1000uF-Electrolytic-Capacitors-11X18/dp/B07GCV44GM/ref=asc_df_B07GCV44GM/?tag=hyprod-20&linkCode=df0&hvadid=241955606060&hvpos=&hvnetw=g&hvrand=15783515294660329827&hvpone=&hvptwo=&hvqmt=&hvdev=c&hvdvcmdl=&hvlocint=&hvlocphy=9053030&hvtargid=pla-669997511473&psc=1>

Rasberry Pi light sensor TSL2591

<https://www.amazon.com/Adafruit-TSL2591-Dynamic-Digital-ADA1980/dp/B00XW2OFWW/ref=sr_1_2?keywords=tsl2591&qid=1583356900&s=electronics&sr=1-2>
