## Appendix 2 for "An inexpensive setup for robust activity tracking in small animals: Portable Locomotion Activity Monitor (pLAM)"

**pLAM Workflow**

**1.Mesh cage:** A mesh cage is needed to contain the animal (Fig 1A,Fig 2A). Cage size influences the animal’s scope of motion should scale with animal size. We used 12”X12” small cages for insects around 5-10mm and medium sized 18”X18”30” cages for insects between 10-50 mm (Raising Butterflies, <https://store.raisingbutterflies.org/>). Cage size and orientation should match the camera field of view to maximise the motion detection range. In very large cages, the organism may be too small to accurately resolve, and it may be  better to use multiple detectors. Softer netting mesh cages are best since they reduce wear and tear on the animal and increases longevity  However, if used outdoors, softer cages are more prone to being disturbed by wind and hard meshes can reduce detection of false motion. The motion detection setup is sensitive to abrupt changes in light intensity and distant motion, such as moving branches, and therefore placement of the setup is critical to obtain accurate results. Cages can have a combination of clear and opaque sides, and it is important to orient them or choose opacity to reduce background motion while maximising illumination. Cages with reflective colors are useful to spread light evenly, whereas black cages make object tracking harder.

**2.pLAM Hardware**

**a) Raspberry Pi and camera:** Raspberry Pis (Fig 1A,2B)  are compact and mobile single board computers that are a fraction of the cost of a laptop or desktop, yet can perform many of the same tasks. Raspberry Pis can easily interface with hardware to control lights, sensors and cameras. They can also interface with software to selectively capture data and perform analysis. The hardware is standardised and easily available through commercial vendors and the software can be shared as an out of the box ready to use disk image that reduces version control issues and software setup, a common barrier for using open-source software.  There are several models Raspberry Pi3 and RaspberryPi 4, but we have optimised pLAM for Pi3 since it is cheaper and has more stable performance. To keep costs low and improve latency, we utilise the Adafruit Raspberry Pi NOIR camera. They are optimised for use with the PI and come with two small inbuilt IR lights that are powered directly by the PI. They also have an IR filter that automatically gets cut when the light is bright to prevent images from getting too washed out during the day. It is possible to use more advanced USB webcams, but they are usually more bulky and expensive, and require their own external power source.

**b) Power bank/ Wall outlet:** The Raspberry Pi’s require a 5V, 2.5 A standard USB input. They can be run with power banks or a wall outlet (mains). It is recommended to run them off mains however, they can be powered by power banks (Fig 1 A, 2A) or larger capacity batteries. The duration of the experiment will determine the size and of the battery needed. Field use tests show they last for 12 hours on a 20,000 mAh, 4.5 V (100 Wh) power bank. For running longer experiments, portable 750WH UPS can be used or very high capacity power banks (50,000mAH) can be used, although airline transport restrictions on larger capacity lithium batteries should be considered. If access to mains are available but unreliable, lower capacity UPS’s or piHAT UPS’s can be used to ensure continuous operation over the trial window. (See Supplementary material)

**c) IR lights:** Many insects and animals alter their behaviour in the presence of visible and UV artificial light ^24,25^ and therefore the setup relies on infrared (IR) illumination in low-light conditions because most insects are unable to see IR lights ^26^. While the Arducam NOIR camera already has two small IR illuminators (Fig 1A, 2B) , we recommend adding extra IR illumination  (Fig 1A, 2A) to the setup to improve performance. Since these require additional power, this may not always be feasible and to record behaviour without these lights, it is better to use smaller mesh cages. As cage size increases and animals become smaller, brighter illumination is required. Several IR light options exist and they vary in cost, power requirements and brightness (See Supplementary Information).

**3. Animal density:** The activity detector detects motion based on differences in pixel values over successive frames. Such values change whenever an animal moves significantly in the field of view of the camera. The threshold of pixels required for a motion event can be modified, but generally smaller animals need to be closer to the camera to cover enough pixels to trigger an event. This holds both for single or multiple individuals but the chance of picking up an event is higher with a higher density of individuals. Higher densities increase the chance of capturing a motion event, but with larger animals, high densities can cause flying animals to damage each other. Thus, it is advised to use higher density of animals if they are small and lower density if they are large.

**4. pLAM Software**

**a) pLAM OS:** The raspberry pi OS we chose for the device is Rasbian, a linux based OS. We provide a disk image of the associated software and packages that can be written to a memory card and be used with minimal updating. For more advanced users we provide the python based code on github (www.github.com/yashsondhi/diel-light-pi) as well as detailed instructions on setting up the associated software from scratch.

**b) Pi setup and headless mode:** The Raspberry Pi usually requires an external monitor, keyboard and mouse to initially configure. These peripherals can be used to start the experiment and then disconnected safely. Alternatively, if the Pi’s are connected to a network, they can be booted up headless (without a monitor, keyboard or mouse) and controlled remotely via ssh (CUI) or VNC Server (GUI). It is recommended to run them on a network with the internet access as this ensures updated system time and lets them send the information required to remotely access the Pi’s on boot (Supporting Information)

**c) pLAM configuration:** The software behind pLAM is Motion, an open-source tool that is widely used for DIY motion tracking projects and has a large community of users. Running and configuring Motion on the Raspberry Pi can be challenging without sufficient programming knowledge. To simplify it, we pre-install Motion on the pLAM OS disk image and created a set of pre-configured Motion configuration files that work for different activity chamber requirements such as animal size, time lapse, video recording and snapshots of each motion event. The user inputs details about the location, organism and choice of these pre-configured settings in a small configuration file. We added features useful for running in the field such as manually checking and updating system time in the absence of an internet connection, auto resume on reboot and providing instructions on using them.

**d) Data transfer and memory:** Monitoring activity using images or videos generates large amounts of data. Although data transfer from the Pi’s can be achieved remotely, as file volumes get larger than a few GB, data transfer becomes a bottleneck. We recommend using larger SD cards and also using the “best” mode wherein the image with the most motion is saved from each motion event. This allows post-facto cleanup and troubleshooting, without generating excess data. A log file for each motion event is generated which can be used for downstream analyses after confirming the validity of the motion event capture using the event images.

**e) Analysis:** We designed scripts to allow analysis along with data generation with the Pis. The scripts for analysis are available on the pLAM OS and can be run on a local computer as well. We allow the user to export a text file with the duration of each event and the pixel difference as a proxy for the amount of motion. These can be plotted as (i) daily activity histograms with a cumulative count. (ii) pixel difference of each event with time. (iii) Frequency probability distribution of activity event durations and (iv) Mean activity across the a 24 hour duration (See supplementary information for more details about each plot)

We include in the supplementary methods a detailed instructable/ guide on setting up your own pLAM, controlling the light environment along with collecting and analysing data (Supplementary information)

**Guidelines for recording diel-activity with pLAM:**
pLAM works consistently well in an isolated lab environment and the same RasberryPi that monitors motion can be used to control ambient lighting. We found that isolating the setup in incubators is ideal, but any closed chamber that is shielded from external light and wind works well. While the Pi can be used to monitor temperature, light and humidity, it is often easier to use commercial bluetooth or wireless loggers to monitor the chambers as they provide accurate data and make the setup easier. It is easier to get the pLAM and lights working in a lab environment at first and the data is much cleaner. Assuming the camera and light system work in isolation, the next factor that is important is the duration of the experiment and the density of animals. For mosquitos, flies and moths, we found that by keeping humidity levels high and providing a food source, we could get data for 5-7 days post eclosion. We found that when using small cages, 20-25 flies or mosquitoes generated sufficient data to give an accurate representation of the activity, but with moths 4-8 individuals was sufficient. It is possible to record even a single individual, but since the animal is not always in the field of view of the camera, it is possible that activity data is incomplete. We found it was useful to limit the insect motion to smaller chambers entirely within the camera's field of view, but this reduces the animal’s tendency to fly, a behaviour we were interested in, so we did not pursue that option further. We noticed for flies at least, we were much more likely to pick up flying behaviour than walking behaviour, but this was not an issue for the larger moths. While setting up pLAM experiments with different organisms, it is likely that several parameters will have to be tweaked and we recommend altering animal density, cage size, using higher resolution cameras, lowering the motion threshold, and using stronger IR illumination to improve the quality of data being generated.
Conducting pLAM trials in the field is slightly more challenging. Access to electrical mains, internet and shelter from wind and rain and light pollution are ideal, but often hard to obtain. Portable routers, portable UPS’s and lightweight tents are relatively inexpensive solutions to some of these problems, but if possible it is advisable to choose periods without heavy wind and rain as they can make data collection very noisy. The field trials were held under harsh weather conditions more prone to interruptions from power failure, random background motion of the forest due to the wind and water leakage on the electrical equipment. Two setups even got blown away by the wind, and we eventually used a large indoor library with windows to provide access to light while shielding the setups from wind. Despite this we were able to collect data for over 10 species in under 10 days and we learnt from our failures from the field to improve the software and mentioned the hardware solutions that improve the robustness of the setup in field-conditions.
