## Supplementary Methods for "An inexpensive setup for robust activity tracking in small animals: Portable Locomotion Activity Monitor (pLAM)"

**a) Animal Care:** Moth Rearing: Wax moths (*Galleria mellonella)* were reared in a temperature controlled incubator at 22°C with 50-60% relative humidity with a gradual day night cycle (14L:10D). Dusk and dawn were set to duration of two hours and gradually increased (Fig. S3), with constant brightness for the day (100-300 lux) and no visible light at night. Moths were ordered from Carolina Biological as final instars and were transferred to medium sized upright plastic containers with holes in their lid to allow some respiration. Moisture built on the walls of the containers and had to be removed often to prevent mold build-up develop. When mold did build up it was only the outer layer of the food source and did not seem to affect the emergence of the moths. Fruit flies (*Drosophila melanogaster)* were reared from an ongoing lab colony, and were fed standard media and reared at 21 °C on a 12 h:12 h light: dark cycle. Larvae continued into adulthood in the same manner. During their third instar and prior to the wandering stage, larvae were separated from the media using a sieve and running water and were placed in a jar with moisture but without media.

**b) Simulating Light cycle and extra IR lighting:** We used a 30 LED strip (Adafruit Neopixel) where we could control the intensity and color of each LED to simulate dusk, dawn and day and night lighting. To cross check the light cycles, we measured the relative light intensity in the chamber lux using the Adafruit TSl2591 sensor. Although the readings were sensitive to changes in position of the sensor they were reliable to measure the relative changes in intensity over the course of the trial. We used 9-12 light IR floodlights (Jcheng Security Infrared illuminator) to film the behaviour of the moths, these lights turned on and off automatically based on the ambient light levels.

**c) pLAM methods:**

*Bead Trials*: We used three different sized beads of dimeter(7mm, 10 mm and 14 mm) and suspended them from the roof of a small wooden cabinet (750 mm high). Beads were suspended at 550, 600 and 650 mm from the pLAM camera and we measured the motion for 10 minutes, 5 minutes light and 5 minutes dark. During the light condition, the room lights on durin the dark condition, visible lights were turned off and the illumination was provided by the IR illuminator. A small portable fan was turned on randomly to simulate motion over the course of the trials.

*Insect trials*: We monitored activity using our setup for different animals according to size. Medium sized mesh cages (13” x 13” x 24” inches) were used with 4-8 moths taken 0-3 days after eclosion. Small sized mesh cages (12” x 12” x 12” inches were used with 20-25 flies taken 1-2 days after eclosion and 20-25 mosquitoes 1-4 days after eclosion. Flies and mosquitoes were kept as a food source of a 7% sugar solution in  a small glass vial. Wax moths are not known to require food as adults and often got stuck in the water and drowned, so we did not place food in the chamber.

 Trials were conducted between 20-22℃ with ambient humidity between 50-60 %. We used a combination of males and females for all species to better recreate natural activity. (Except mosquitoes, where only males were used). We monitored the cumulative behaviour using the activity detector. We captured every motion event and saved an image with the maximal motion and used this as a proxy for activity.

**d) LAM methods:** To compare our methods to previously published results cite) we used moths and flies in this device. Trials were conducted in a temperature controlled incubator at 27℃. Moths were kept in a chamber of dimensions 8X8X2 inches with each cube of size  2.3X2.3X 2 inches. A 30 LED strip was used to illuminate the chamber.  We placed 8 males and 8 females 2 days after eclosion in the LAM tubes with a food source consisting of x percent sucrose solution. We placed 32 flies in an LAM and they were kept for 5 days in the same conditions but instead of adults, pupae close to eclosion were used. 16 Male *Aedes aegypti* were tested over the course of 5 days (supplementary)

**e) Field Methods**
We conducted field trials at the Estación Biológica Monteverde (EBM), Puntarenas Province, Monteverde, Costa Rica (1530 m, 10°19'08.5"N, 84°48'32.0"W) from 1-10 July 2021. EBM is a private research station within a cloud forest preserve, and is located on the Pacific slope of the Tilarian Mountain Range. We set up light traps with 250W metal halide (6500K) lamps and LepiLed (https://www.gunnarbrehm.de/en/lepi-led) lights at multiple locations around the station. Species that we could identify easily in the field were collected and tested in the pLAMs, although some species were only identified to the genus and family level. We used 6 pLAM detectors with IR illuminators and ran each trial for ca. 48 hours. Each pLAM had between 4-12 individuals of the same species and we digitised each individual after completion of the trial using a Nikon D7500 with a Nikkor 105mm macro lens. We conducted four sets of trials. Trial #1 was conducted outdoors in tents that were open on one side to ensure natural light. Weather conditions worsened by 5th July 2021 and the strong wind and rain disrupted further trials (#2). We moved the pLAM’s indoors in a room with sufficient natural light for the remaining trials (#3-4). (Table S4)
